# PKA prevents resection of DNA double strand breaks and favor non-homologous end-joining

**DOI:** 10.1101/2024.08.29.610313

**Authors:** Houaria Khater-Sellou, Indiana Magdalou, Vasily Ogryzko, Bernard S. Lopez

## Abstract

Genome rearrangement is a hallmark of cancer and ageing. DNA double-strand breaks (DSBs) are highly toxic lesions that can generate genome rearrangements. Several pathways compete for DSB repair, and selection of the appropriate repair process is decisive for genetic stability. DSB repair act according to two steps: 1-the choice between nonhomologous end-joining (NHEJ) *versus* resection of DNA ends, generating single-stranded DNA (ssDNA); 2-on ssDNA, homologous recombination (HR) and sub-processes. Here we show that 53BP1, which plays a prime role at the first step by protecting against resection and fostering NHEJ, physically interacts with the catalytic subunits of the cAMP dependant protein kinase A (PKAcs). PKA favors the recruitment of 53BP1 at DNA damages sites, and consistently, prevents resection, favoring NHEJ. Inhibition of PKA stimulated resection and coherently reduces NHEJ. Reversely, activation of PKA with 8-Bromo-cAMP stimulate NHEJ and reduces HR. These data open new avenues for potential anticancer strategies. More generally, they underline the high complexity of the regulation of DSB repair, identifying PKA as a novel actor of the DNA damage response, acting at the choice of the DSB repair process, which might be essential for genome integrity maintenance.

## Introduction

Genome instability is a hallmark of both cancer and aging (Negrini et al. 2010; Hanahan and Weinberg 2011; López-Otín et al. 2013; Garinis et al. 2008; Hoeijmakers 2009; López-Otín et al. 2023). Genome is daily assaulted by different stresses from endogenous as well as exogenous stresses. Remarkably, activation of the DNA damage response (DDR), has been observed at early stages of cancer and senescence (Bartkova et al. 2005; Gorgoulis et al. 2005; Bartkova et al. 2006), suggesting that genetic stresses represents an initial step in these processes.

The DNA double-strand break (DSB) is considered as the most toxic lesion, and is also a prominent source of genetic instability through both genome rearrangements and mutagenesis at the scares of the repair junctions. However, DSB repair is also employed by the cell to generate profitable genetic diversity in physiological processes such as the establishment of the immune repertoire, meiosis and neuronal gene expression (for review, see So et al. 2017). In these physiological situations, DSBs are generated by controlled cellular enzymes then cells employed the same DSB repair machineries than that for accidental DSBs. Consequently, DSB repair should be finely regulated to maintain the balance between genetic stability *versus* diversity. Therefore, the selection of the most adapted DSB repair pathway becomes thus a crucial issue.

Two main strategies are employed to repair DSBs: the first one is named nonhomologous end-joining (NHEJ) and ligates two DNA double-strand ends without requiring sequence homology. In this process the Ku70/Ku80 heterodimer binds to the DSB and recruits DNA-PKcs, Ligase IV and its cofactors such as XRCC4 and Cernunos/XLF (Guirouilh-Barbat et al. 2004, 2007; Bétermier et al. 2014; Deriano and Roth 2013; So et al. 2017; Scully et al. 2019).

The second DSB repair strategy is based on the use of homologous sequences borne by an intact DNA molecule that will be copied; this mechanism is thus called homologous recombination (HR). HR acts on 3’-single-stranded DNA (ssDNA) arising from the resection of the DSB ends, on which RAD51 is loaded by BRCA2. This generates a ssDNA/RAD51 filament that promotes the search for the homologous partner and strand invasion.

The choice of the DSB repair process operates in two steps (Bétermier et al. 2014; Rass et al. 2009; So et al. 2017, 2022): 1-the selection of NHEJ *versus* resection; 2-on resected ssDNA, HR and sub-processes. Consistent with this model, studies in mammalian cells have shown that defects in NHEJ stimulate HR (Guirouilh-Barbat et al. 2007; Delacôte et al. 2002; Pierce et al. 2001; Guirouilh-Barbat et al. 2004). 53BP1 plays a prime role at this initial step. Indeed, 53BP1 protects DNA ends from resection, thus favoring NHEJ, and depletion of 53BP1 abrogates HR defects of BRCA1-deficient human cells (Bunting et al. 2010).

NHEJ is efficient all along the cell cycle, while HR acts in the S/G2 phases. Therefore, they compete in late S and G2 phases. Thus, in order to deepen the impact of 53BP1 on the choice of the DSB repair mechanism, we performed a proteomic screen by immunoprecipitation of 53BP1 in G2 phase and mass spectrometry analysis, in order to identify partners of 53BP1 that might participate to the choice of the DSB repair process. This revealed the association of 53BP1 with the cAMP-dependent protein kinase catalytic subunit alpha (PKAcs).

PKA has been described to act on DNA-PKcs, Ligase IV, XRCC4 and Cernunos/XLF, thus at late steps, downstream the early step of choice between NHEJ and resection, but contradictory consequences on NHEJ efficiencies have been reported (Noh and Juhnn 2020; Jessulat et al. 2021; Yang et al. 2016; Castejón-Griñán et al. 2018; Huston et al. 2008; Zhang and Steinle 2013) Here we show that PKAcs physically interacts with 53BP1 and favors the recruitment of 53BP1 on damaged DNA following ionizing radiation (IR). Consistently, we show that PKA protects against resection, and favors NHEJ. These data reveal that PKA acts also at the early steps of DSB repair. They open new avenues for potential anticancer strategies, and they underlined the high complexity of the regulation of DSB repair, identifying a novel actor, PKA, as an actor of DDR, at the choice of the DSB repair process, which might be essential for genome stability maintenance.

## Material and Methods

### Cells

GC92 (Rass et al. 2009) and RG37 (Dumay et al. 2006) cells are derivated from a common SV40 -transformed fibroblast cell line, a gift from R.J. Monnat Jr (University of Washington). They were originally obtained from the National Institute of General Medical Sciences Human Genetic Mutant Cell Repository (Camden, N.J.). CG92 cells contain the CD4-3200bp substrate, which monitors the EJ-mediated deletion of a 3200bp fragment by expression of the membrane antigens CD4 (Guirouilh-Barbat et al. 2004; Rass et al. 2009; Gelot et al. 2016; Guirouilh-Barbat et al. 2007; Grabarz et al. 2013; Guirouilh-Barbat et al. 2016a; So et al. 2022). RG37 contain the DR-GFP substrate (Pierce et al. 1999) that monitor HR through GFP expression.

AID-DIvA (AID-AsiSI-ER-U20S) is a U2OS cell line (human osteosarcoma) in which the AsiSI endonuclease induced DSBs at specific regions in the genome AsiSI is sequestered in the cytoplasm, but addition of 4-hydroxy-tamoxifen (4OHT; 300 nM, Sigma-Aldrich, H7904) to the culture medium for 4 h, induce the translocation of AsiSI into the nucleus, which then cleaves DNA. After 4OHT treatment, cells are washed three times in prewarmed PBS and further incubated with 500 µg/ml auxin (IAA) (Sigma-Aldrich, I5148) for 2 h to induce the degradation of AsiSI (Aymard et al. 2014). Then, the medium is refreshed, and the cells are cultured for 18-20 h before DNA collection.

All cell lines were cultured in DMEM supplemented with 10% FCS and were checked monthly for mycoplasma contamination by PCR (primers 5’ GGGAGCAAACAGGATTAGATACCCT 3’ and 5’ TGCACCATCTGTCACTCTGTTAACCTC 3’)

### Transfection

The meganuclease I-SceI, was expressed by transient transfection of HA-I-SceI expression plasmid (Liang et al. 1998) in RG37 and GC92 cells with Jet-PEI following the manufacturer’s instructions (Ozyme, 101-40N). The expression of HA-tagged I-SceI was verified by Western blotting. For RNA silencing experiments, 25,000 cells were seeded 1 day before transfection performed using INTERFERin following the manufacturer’s instructions (Polyplus Transfection, New York, NY, USA, #409-50) with 20 nM of one of the following siRNAs: Control (5’ UAAGGCUAUGAAGAGAUAC(dTdT) 3’), synthesized by Eurofins (Ebersberg, Germany)), PKAcs (Cell signaling, 6574). Seventy-two hours later, the cells were transfected with the pCMV-HA-I-SceI expression plasmid.

### Immunofluorescence

Immunofluorescence experiments were performed on cells grown on glass coverslips. Cells were fixed with 4% paraformaldehyde and were then permeabilized with 0.2% Triton X-100 for 15 min at RT. Soluble proteins were extracted before fixation by incubating coverslips with extraction buffer (50 mM Tris HCl (pH 7.4), 150 mM NaCl, 1% Triton X-100 and protease inhibitor cocktail (cOmplete Mini Protease Inhibitor, Roche, 5892970001) for 5 min on ice. After blocking in PBS containing 3% BSA and 0.05% Tween 20, immunostaining was performed using mouse anti-53BP1 (1:500, Merck millipore, MAB3802), rabbit anti-PKAc (1:100, Cell Signaling, mAb 5842) After 2 washes with PBS containing 0.05% Tween 20, the coverslips were incubated for 30 min with Alexa Fluor 488- and/or 568-conjugated anti-mouse (Life Technologies,1/500, A11029) and anti-rabbit secondary antibodies (Life Technologies,1/1000, A11036). All incubations were performed for 45 min at RT with antibodies diluted in PBS containing 3% BSA and 0.05% Tween 20. After 2 washes, the coverslips were mounted in mounting medium (Dako, S302380-2) supplemented with DAPI (Sigma-Aldrich). Images were acquired using a Leica SPE confocal laser scanning microscope or an Olympus BX63 microscope with a 63× oil objective. Images were imported, processed and merged in ImageJ software.

### Analysis of DSB repair at the CD4-3200bp intrachromosomal reporters

After transfection with the I-SceI expression plasmid and incubation for 72 h, cells were collected with 50 mM EDTA diluted in PBS, pelleted and fixed with 2% paraformaldehyde for 10 min. The cells were incubated for 10 min with 1 µl of an Alexa Fluor 647-conjugated anti-CD4 antibody (rat isotype, RM4-5, Invitrogen). The percentages of CD4-expressing cells were determined by FACS analysis using a BD Accuri C6 flow cytometer (Becton Dickinson). To eliminate variability due to the transfection efficiency, all values were normalized to those for control cells transfected with the I-SceI plasmid alone.

Where indicated, cells were treated with 20 µM PKA inhibitor H89 (Sellekchem, S1582), Bromo-8-cAMP 50µM PKA activator (Sigma, B5386), during the first 48 h after I-SceI transfection. Then, the medium was refreshed for the remaining 48 h of incubation before cells were collected for FACS analysis.

#### Western blot analysis

Cells were lysed in buffer containing 50 mM Tris HCl (pH 7.5), 20 mM NaCl, 1 mM MgCl_2_, and 0.1% SDS supplemented with Complete Mini Protease Inhibitor (Roche) and treated with 250 U of benzonase (Santa Cruz, sc202391) for 30 min. Proteins (30–40 µg) were denatured, separated on 9% SDS‒PAGE gels, and transferred onto nitrocellulose membranes, which were incubated with the following specific antibodies: rabbit anti-PKA (1:500, Cell Signaling, mAb 5842, mouse anti-HA (1:1500, Santa Cruz, sc-7392), rabbit anti-pCREB (1:1000, Cell Signaling,♯9198), anti-53BP1 (1:100, Merck millipore, MAB3802) and mouse anti-Vinculin (1:5000, Abcam, ab18058). Immunoreactivity was visualized using an enhanced chemiluminescence (ECL) detection kit (Pierce).

#### Proximity ligation assay (PLA)

Cells grown on coverslips were fixed with 2% paraformaldehyde for 10 min, permeabilized, blocked and prepared as described above for immunostaining with the following primary antibody pairs: rabbit anti-53BP1 (1:100, Cell Signaling, ♯4937) and mouse anti-PKAcs (1/100, cell signaling, mAb 5842).

PLA was performed using a Duolink In Situ Detection Kit (Sigma-Aldrich, DUO92001, DUO92005, DUO92008) according to the manufacturer’s protocol. Images were acquired with a Leica SPE confocal laser scanning microscope using a 63× objective lens. Images were processed with ImageJ software.

#### Coimmunoprecipitation

Cellular proteins were extracted on ice using 25 mM Tris–HCl (pH 7.5), 150 mM NaCl, 1 mM EDTA, 0.5% NP40 and cOmplete Mini Protease Inhibitor (Roche). Protein extracts were treated with DNase I (15 U/ml, Thermo Scientific, EN0521) for 30 min at RT. Extracts were precleared with Dynabeads (Life Technologies, 10004D) for 30 min at 4 °C, and 300 µg of protein was then incubated with 1 µg of a mouse anti-53BP1 antibody (1μg/ml, Merck millipore, MAB3802) O/N at 4 °C. Then, 25 µl of Dynabeads was added, and the mixture was incubated for 4 h at 4 °C. The beads were subsequently washed three times with extraction buffer. Laemmli buffer (2×) with 4% ß-mercaptoethanol was used to dissociate and denature the bead-antibody-protein complexes. Western blot analysis was performed to detect PKAcs and 53BP1 using a rabbit anti-53BP1 antibody (1:1000, Cell Signaling, ♯4938) and a rabbit anti-PKAcs antibody (1:500, Cell Signaling, mAb 5842), respectively.

### Foci counts

Foci were automatically counted with ImageJ using the following method.

To define a mask with nuclei in the DAPI channel, Image/Adjust/Threshold/Analyze/Analyze Particles was used. In the foci channel, all particles in the ROI manager were selected, followed by “OR (Combine)”. All nuclei are outlined in the foci image. To define the threshold for counting foci, we used Process/Find Maxim and “Single Points” as the output type, and we determined the correct value for detecting the majority of foci (this value should not vary between images in the same experiment). A new window then appeared. In the ROI Manager, “Measure” was used. In the results datasheet, the foci number by nucleus was obtained by dividing the “Raw Integrated Density number” value by the “Max” value (255).

### Resection assay

Resection measurements on DIvA cells were performed as previously described (Zhou et al. 2013). After 4-hydroxy-tamoxifen treatment, cells were collected for genomic DNA extraction (DNeasy blood & tissue kit, Qiagen, Hilden, Germany), and 100-200 ng genomic DNA was treated with 16 U of the appropriate restriction enzyme overnight at 37°C. After enzyme inactivation, the digested genomic DNA was used for qPCR (mix for qPCR, TAKARA, Shiga, Japan) with the primers listed in the table below.

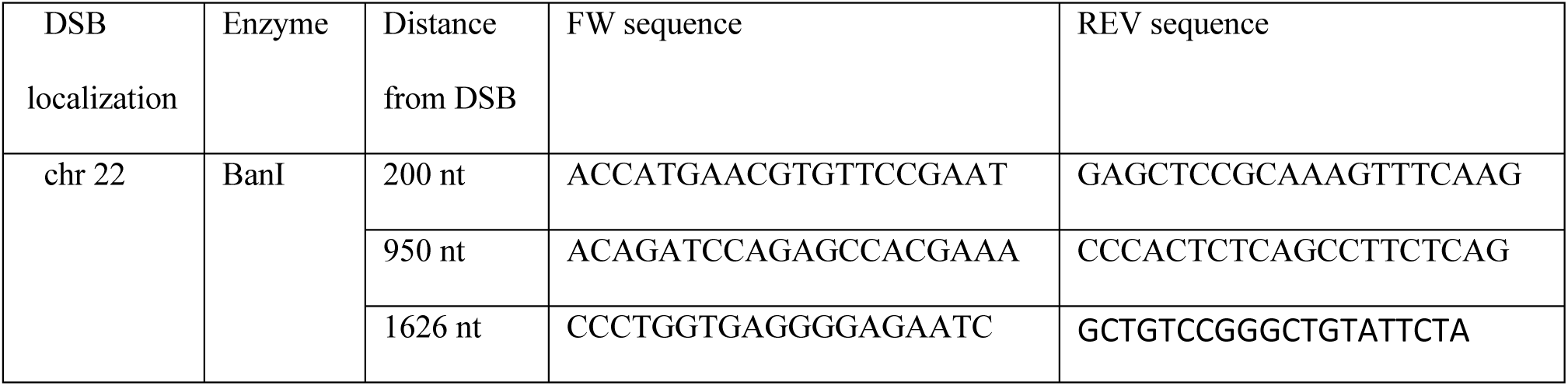

The percentage of ssDNA was calculated with the following equation: ssDNA % = 1/(2^(△Ct-1) + 0.5)*100, where △Ct = Ct of the digested sample – Ct of the nondigested sample.

#### Synchronization in G2 with RO-3302

Cells were treated with Le 5 µM RO-3306 (Calbiochem #217699) before analysis 20 h avec le RO-3306 avant d’être fixées.

#### nLC-MS/MS analysis

Proteins were extracted with 190 mM Amonium bicarbonate and digested overnight using a on-beads digestion method with trypsin at 37°C at pH 7.8. Eluted peptides were desalted on Vivapure C18 micro spin columns (Sartorius Stedim Biotech), desiccated in SpeedVac.

Dried peptides were resuspended in 10µl of 0.1% formic acid and 1µl was injected for nLC-MS/MS analysis, in triplicate. Liquid chromatography and mass spectrometry analyses were performed on an EASY-nLC 1000 paired with Q Exactive quadrupole-orbitrap hybrid mass spectrometer (both from Thermo Fisher Scientific). The peptides of each sample were separated on EASY-Spray 15 cm × 75 µm 3 µm 100Å C18 PepMap® reverse-phase column (Thermo Fisher Scientific) using 150 min three-step water-acetonitrile gradient (0-120 min, 5 → 35% LC buffer B (0.1% formic acid in acetonitrile); 120-140 min, 35 → 50%; 140-145 min, 50 → 90%; hold for 5 min) at 300 nL/min flow rate. The mass spectrometers acquired data in positive mode throughout the elution process and operated in a data-dependent scheme with full MS scans acquired, followed by up to 10 successive MS/MS fragmentations and measurement on the most abundant ions detected. The MS scan range was set up at 400-2000 m/z, with a resolution of 70,000, an AGC target of 1E6 and e MIT of 100 ms. MS2 measurement was set up at a resolution of 17,500, an AGC target of 5E4, a MIT of 100 ms and an isolation window of 2 m/z, a collision energy of 27 and with 30 sec dynamic exclusion.

Raw mass spectrometric data were analyzed by Proteome Discoverer using the Homo Sapiens Swissprot database. A maximum of 2 missed cleavages was allowed. The precursor mass tolerance was set to 10 ppm and the fragment mass tolerance to 0.02 Da. Carbamidomethylation of cysteins was set as constant modification Deamidation of asparagine and glutamine and oxidation of methionines were set as variable modifications. False discovery rate (FDR) was calculated using Percolator with 0.01 strict and 0.05 relaxed target cut-off values.

### Statistical analysis

Unpaired t-tests were performed using GraphPad Prism 3.0 (GraphPad Software).

## Results

### PKAcα interacts with 53BP1

We have analyzed by mass spectrometry the interactors of 53BP1 in cells synchronized in G2 with RO-3306 through co-immunoprecipitation with 53BPA antibodies (data to be published). The relevance of these interactions was attested by the fact that the canonical interactors of 53BP1, i.e. Tp53, and DNA-PKcs, which plays an important role in NHEJ, were both identified in the proteomic analysis.

Remarkably, the data also revealed the presence of PKAcα. Additionally, cAMP-dependent protein kinase regulatory type II-α and type II-β (RIIα and RIIβ) subunits were also identified, supporting an interaction between 53BP1 and PKA type II.

Then we tested the physical interaction between PKAcs and 53BP1 by co-immunoprecipitation in a human cell lines (SV40 transformed fibroblasts). In asynchronous cells, immunoprecipitation of 53BP3 did not significantly pull down PKAcs (Figure 1A). However, in G2 synchronized cells (with RO-3306), PKAcs, efficiently co-immunoprecipitated with 53BP1 (Figure 1A). To confirm these data and the sub-cellular localization, with performed a proximity ligation assay (PLA) (Figure 1B). The data confirmed the close colocalization of PKAcs, and 53BP1 in G2 synchronized cells but not in asynchronous cells (Figure 1B). In addition, PLA reveal the co-location of PKAcs, and 53BP1 into the nucleus. Collectively, these data show the interaction of PKAcs and 53BP1 into the nucleus in G2 arrested cells.

**Figure 1.**
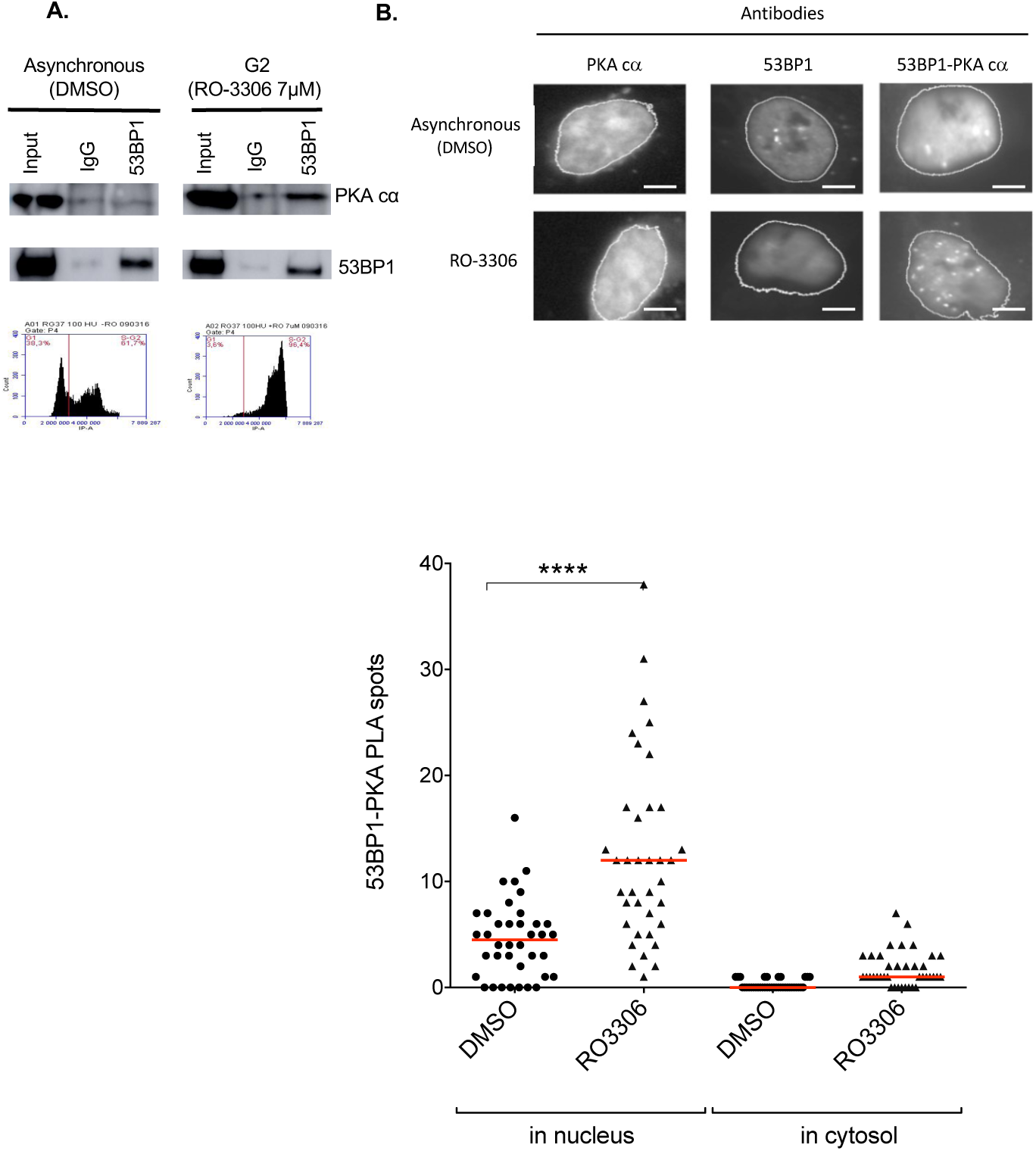
Physical interaction between 53BP1 and PKA in G2/M. **A.** Co-immunoprecipitation in asynchronous (treated with DMSO) cells *versus* G2/M (RO-3306) arrested cells. The lower panel shows the impact of DMSO (left panel) and RO-3306 (right panel) on cell cycle distribution. **B. Proximity ligation assay.** Upper panel: representative images of PLA with PKAcs antiboby alone (left panel), 53BPA antibody alone (middle panel), both PKAcs and 53BP1 antibodies (Right panel). Scale bars : 10 µm. Lower panel: quantification (median) of at least 3 independent experiments (*p* values: *<0.05, **<0.01).

Following exposure to a genotoxic stress (here hydroxyurea, HU, which generates a replication stress), cells were less efficiently arrested in the G2/M phase than with RO-3306 and more cells were arrested in the S phase, notably at the highest dose, 100 μM HU (compare Figure 1A and Figure 2A). Nevertheless, PKAcs and 53BP1 were also able to interacted in cells arrested treated with HU, in a dose dependent manner (Figure 2).

**Figure 2.**
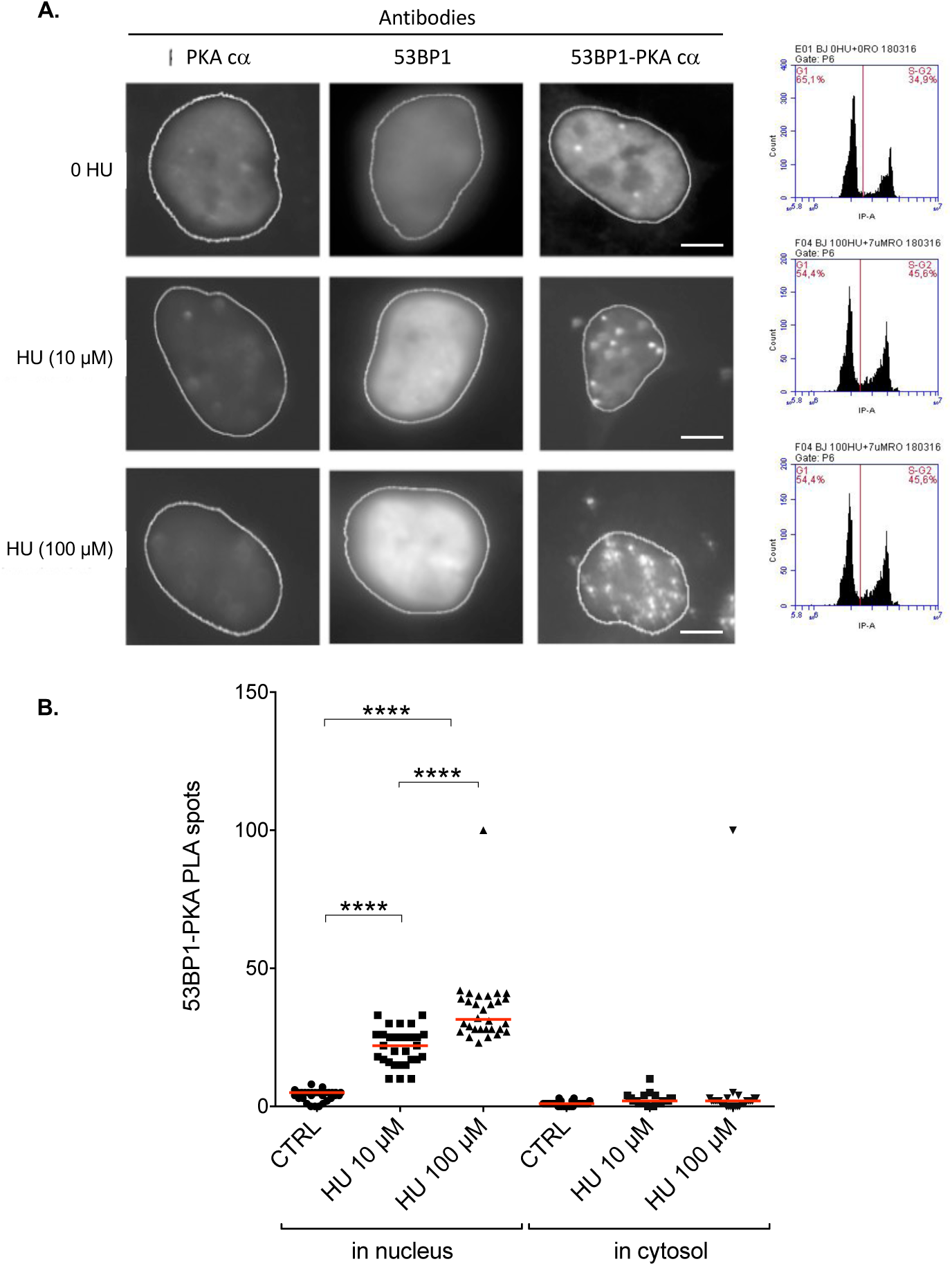
PLA 53P1/PKA after exposure to hydroxyurea. **A.** Representative images of PLA with PKAcs antiboby alone (left panel), 53BPA antibody alone (middle panel), both PKAcs and 53BP1 antibodies (Right panels), at two different doses of HU. Scale bars : 10 µm. Right panels: impact of the treatment on cell cycle distribution. **B.** Quantification (median) of at least 3 independent experiments ( *p* values: *<0.05, **<0.01).

### PKAcs favors the assembly of 53BP1 at DNA damaged sites

53BP1 assembles into nuclear foci, corresponding to DNA damaged sites in chromatin (Anderson et al. 2001). Ionizing radiation (IR) is a convenient genotoxic treatment to measure kinetics of 53BP1 foci formation (Figure 3 A) because the treatment is short (around 2 Gy/min) and then the kinetics are monitored from a precise starting point. The peak of 53BP1 foci assembly is between 30 and 45 min after 6 Gy irradiation (Figure 3B). Remarkably, PKA is activated by IR, as monitored by the phosphorylation of its effector CREB, with a peaked 30 to 45 min. after radiation and this is inhibited by the PKA inhibitor H89 (Figure 3C and 3D), i.e. corresponding to the kinetic of 53BP1 foci assembly. Silencing PKAcs with an siRNA reduced the efficiency of 53BP1 foci formation, 30 to 45 min after IR (Figure 3B). These data show that PKA is an actor of DSB response, favoring the assembly/stabilization of 53BP1 at damaged DNA.

**Figure 3.**
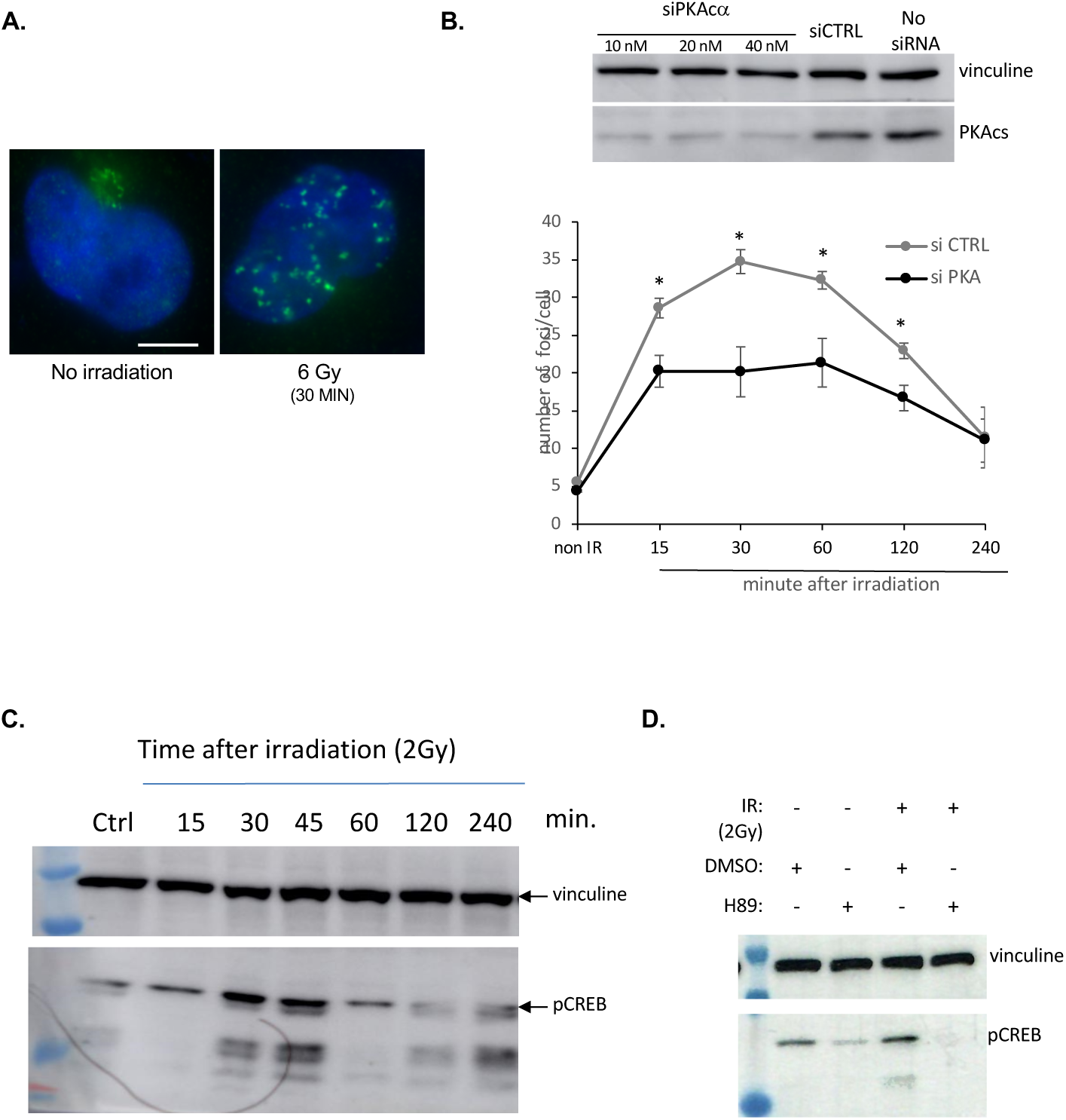
Impact of PKA on 53BP1 foci assembly after ionizing radiation (IR). **A.** Representative picture of nuclear 53BP1 foci after IR (right panel). Scale bar : 10 µm. **B.** Kinetics of 53BP1 foci assembly after IR (2Gy). Upper panel: silencing of PKAcs with siRNA. Lower panel: quantification of 53BP1 (means ± SEMs) foci assembly at different time after IR, with siCTRL (sicontrol) or siPKAcs (20 nM). The data represent at least 3 independent experiments (*p* values: *<0.05). **C.** Kinetics of phosphorylation of pCREB after IR. **D.** Impact of the PKA inhibitor H89 of IR-induced pCREB.

### PKA prevents excessive resection of DSBs

The first step of the selection of the competing DSB repair pathways acts through the choice between resection initiation *versus* protection against resection of the DSB (Bétermier et al. 2014; Rass et al. 2009; So et al. 2017). The absence of resection orientates DSB repair toward NHEJ, while resection engages DSB repair to HR. This choice should be tightly controlled to optimize DSB repair with other cellular parameters as for instance cell cycle; indeed, HR is active only in S/G2 phases while NHEJ is active all along cell cycle (Guirouilh-Barbat et al. 2008; Saintigny et al. 2007; Delacôte et al. 2002; Delacote et al. 2008). Errors on the appropriate DSB repair pathway selection can jeopardize genome stability. By preventing resection, 53BP1 plays a prime role in this choice, and thus favors NHEJ. Therefore, because of the above data we tested the impact of PKA on DSB resection.

We used the DIvA system that allows quantifying the resection at different distances of a Asi-SI cutting site in the genome (Zhou et al. 2013; Cohen et al. 2018). Nuclear translocation of the restriction enzyme Asi-SI was induced by 4-hydroxy-tamoxifen (4-OH-TAM), which then cleaves at restriction sites into the genome, then DNA was extracted. Resection of the Asi-SI-induced DSB will generate single stranded DNA, which is resistant to subsequent *in vitro* cleavage by restriction enzyme. Because the Asi-SI sites have been mapped, we monitored cleavage at different restriction sites located at different distance from the Asi-SI site. Using specific primers, PCR allows to quantify uncleaved DNA (resulting from resection) and thus resection extend (Figure 4A). We measured resection at a site located on chromosomes 22 that has been mapped and used in previous studies (Zhou et al. 2013; Cohen et al. 2018). Resection decreased with the distance from the Asi-S1 cleavage site, as predicted, thus validating the approach (Figure 4B). As a control, silencing CtIP, which initiates resection, significantly decreased the resection efficiency (Figure 4B). In contrast, silencing 53BP1 strongly stimulated resection (Figure 4B), consistently with its role in protection against resection. Coherently with the above data, silencing PKAcα, also stimulated resection. Therefore, PKA favors protection against resection.

**Figure 4.**
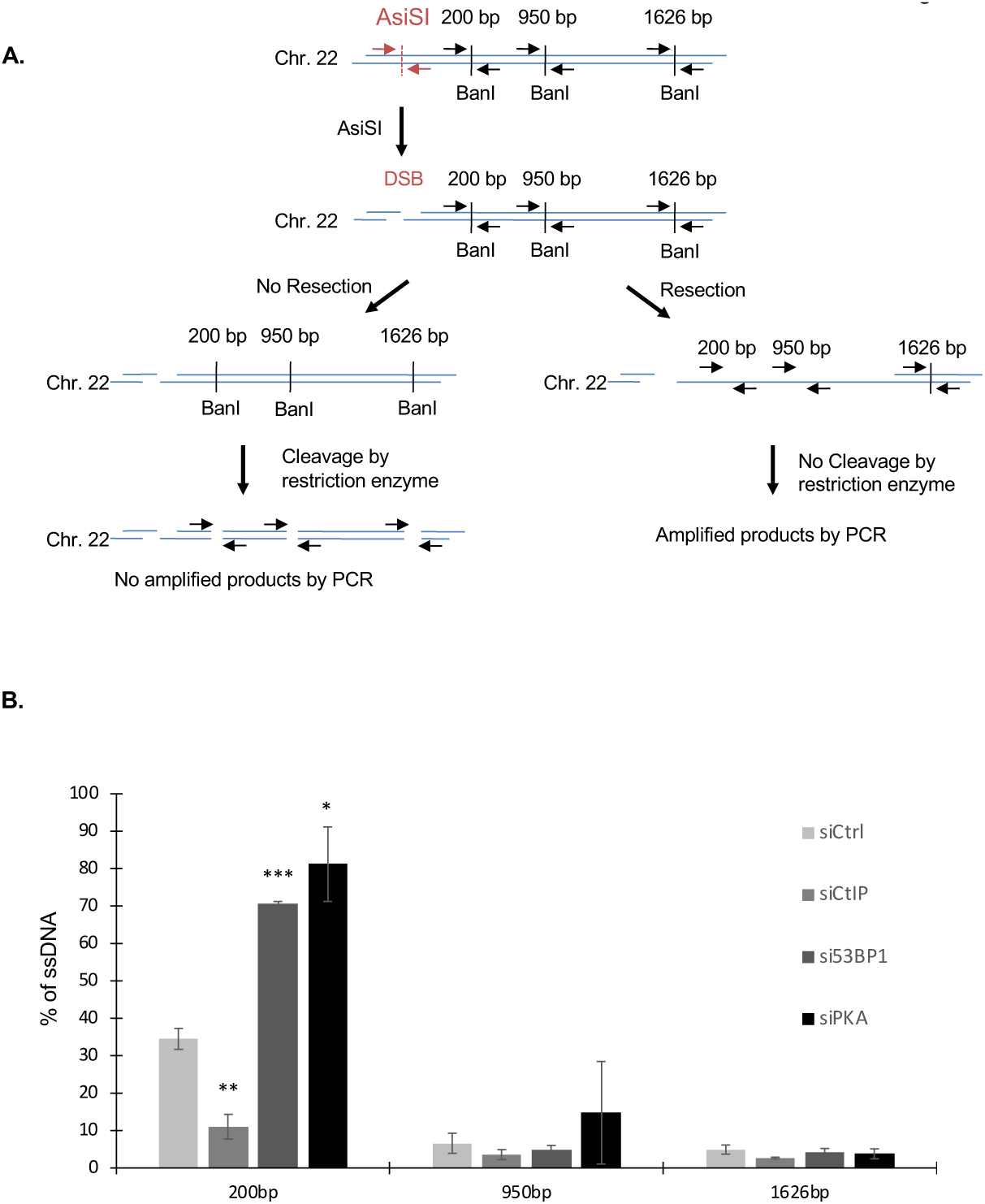
Impact of PKA on resection. **A.** Scheme of the strategy employed. Tthe DIvA system is U2OS cell containing an inducible Asi-SI endonuclease (Zhou et al. 2013; Cohen et al. 2018). Nuclear translocation of the restriction enzyme Asi-SI was induced by 4-hydroxy-tamoxifen (4-OH-TAM), which then cleaves at restriction sites, then DNA. Resection of the Asi-SI-induced DSB will generate single stranded DNA, which is then resistant to subsequent *in vitro* cleavage by the restriction enzyme BanI. We monitored cleavage at different Ban I cleavage restriction sites located at different distance from the Asi-SI site on chromosome 22 that has been mapped and used in previous studies (Zhou et al. 2013; Cohen et al. 2018). Using specific primers, PCR allows to quantify uncleaved DNA (resulting from resection) and thus resection extend. **B.** Quantification of resection different siRNA indicated on the figure. The distance of the BanI from the Asi-SI site are indicated. The quantification corresponds to the means (± SEM) of 5 independent experiments.

### PKA favors NHEJ

Because 53BP1 prevents resection and fosters NHEJ, we then measured the impact of PKA on NHEJ. To monitor NHEJ we used the reporter substrate described in Figure 5A, in human SV40-transformed fibroblast (CG92) and that has been largely characterized (Guirouilh-Barbat et al. 2004, 2007, 2016b; Rass et al. 2009; Grabarz et al. 2013; So et al. 2022; Matos-Rodrigues et al. 2023). Silencing PKAcs with siRNA decreased the efficiency of NHEJ (Figure 5B). Then, we compared the impact of stimulation *versus* inhibition of PKA, using Bromo-8-cAMP or H89, respectively. Inhibition of PKA with H89 confirmed the decrease NHEJ efficiency (Figure 5C). Reciprocally, activation of PKA activity with 8-Bromo-cAMP stimulated NHEJ efficiency (Figure 5C). Moreover, the stimulation of NHEJ by Bromo-8-cAMP was abrogated by co-treatment with H89 (Figure 5B). Collectively, these data show that the activity of PKA favors NHEJ efficiency. These data are consistent with the fact that PKA favors 53BP1 assembly to damaged DNA and prevents resection of double-strand DNA ends.

**Figure 5.**
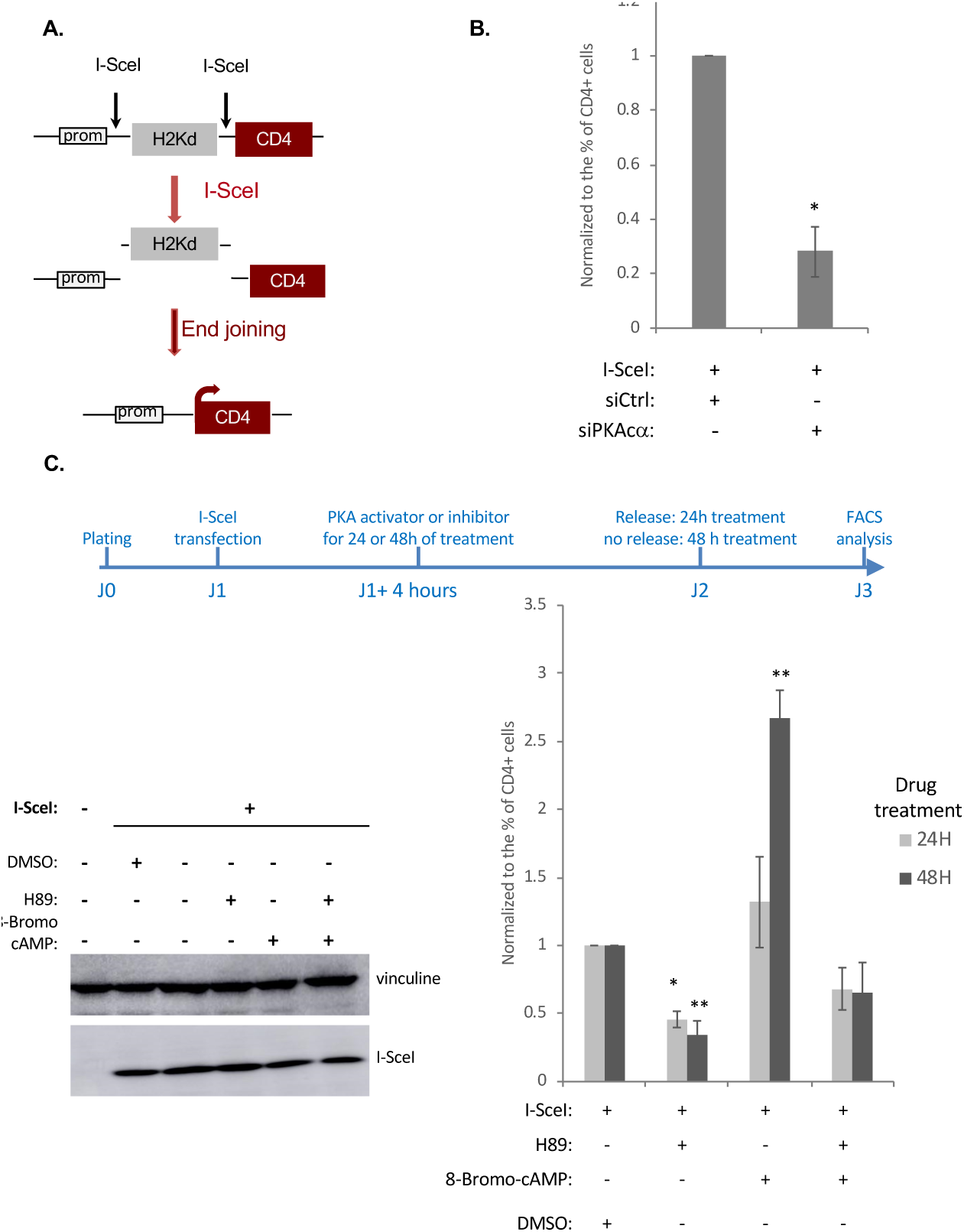
Impact of PKA on NHEJ. **A.** Strategy and reporter substrate used. Here the substrate is chromosomally integrated into the genome of SV40-transformed human fibroblasts (Rass et al. 2009). Before cleave cells express the reporter H2Kd. The reporter CD4 is not expressed because it is too far from the promoter (prom). Two cleavage sites for the meganuclease I-SceI (18 nt recognition) are inserted, one between the promoter and the H2Kd sequence and the other between the H2Ks and CD4. Expression of I-SceI generates the targeted cleavage of the two I-SceI sites and the excision of the H2Kd intervening fragment. End-joining of the DSBs that bear no sequence homology, put the CD4 reporter under the control of the promoter leading to its expression. The frequency of CD4-positive cells gives thus an estimation of NHEJ efficiency. This substrate has been largely validated and characterized (Guirouilh-Barbat et al. 2004, 2007; Rass et al. 2009; Grabarz et al. 2013; Guirouilh-Barbat et al. 2016b; So et al. 2022; Matos-Rodrigues et al. 2023). **B.** Impact of silencing PKA with siRNA. The values are shown normalized to the control siRNA and represent the average ± SEM of 3 independent experiments. **C.** Impact of stimulating PKA (8-Bromo-cAMP) *versus* inhibition (H89) on NHEJ. Upper panel: scheme of the experiment. Lower left panel: expression of I-SceI under the different conditions. Lower Right panel: NHEJ efficiency under the different conditions. The values are shown normalized to the control (DMSO) and represent the average ± SEM of at least 3 independent experiments.

### Stimulation of PKA affect homologous recombination

Then we tested the impact of PKA on HR that is initiated by resection, and that, *in fine,* competes with NHEJ. In a siRNA screen silencing of PKA has been found to stimulate HR (Adamson et al. 2012). This is consistent with the fact that we show here that silencing PKA increases resection (see Figure 4), which corresponds to the first step of HR. Therefore, we addressed the question of the impact of PKA stimulation. To monitor HR we used the DR-GFP substrate (Pierce et al. 1999) (Figure 6A) that is introduced human SV40-transformed fibroblast (RG37) (Dumay et al. 2006), which share the same origin than the GC92 cell line used for monitoring NHEJ above (Rass et al. 2009). Since HR is strictly cell cycle controlled, being active in S/G2 phases. We thus first verified whether stimulation of PKA might affect cell cycle distribution in RG37 cells. Treatment with Bromo-8-cAMP lead to no significant modifications of the cell cycle distribution, depending on the repeat experiment, maybe a small accumulation in the G1 phase (Figure 6B).

**Figure 6.**
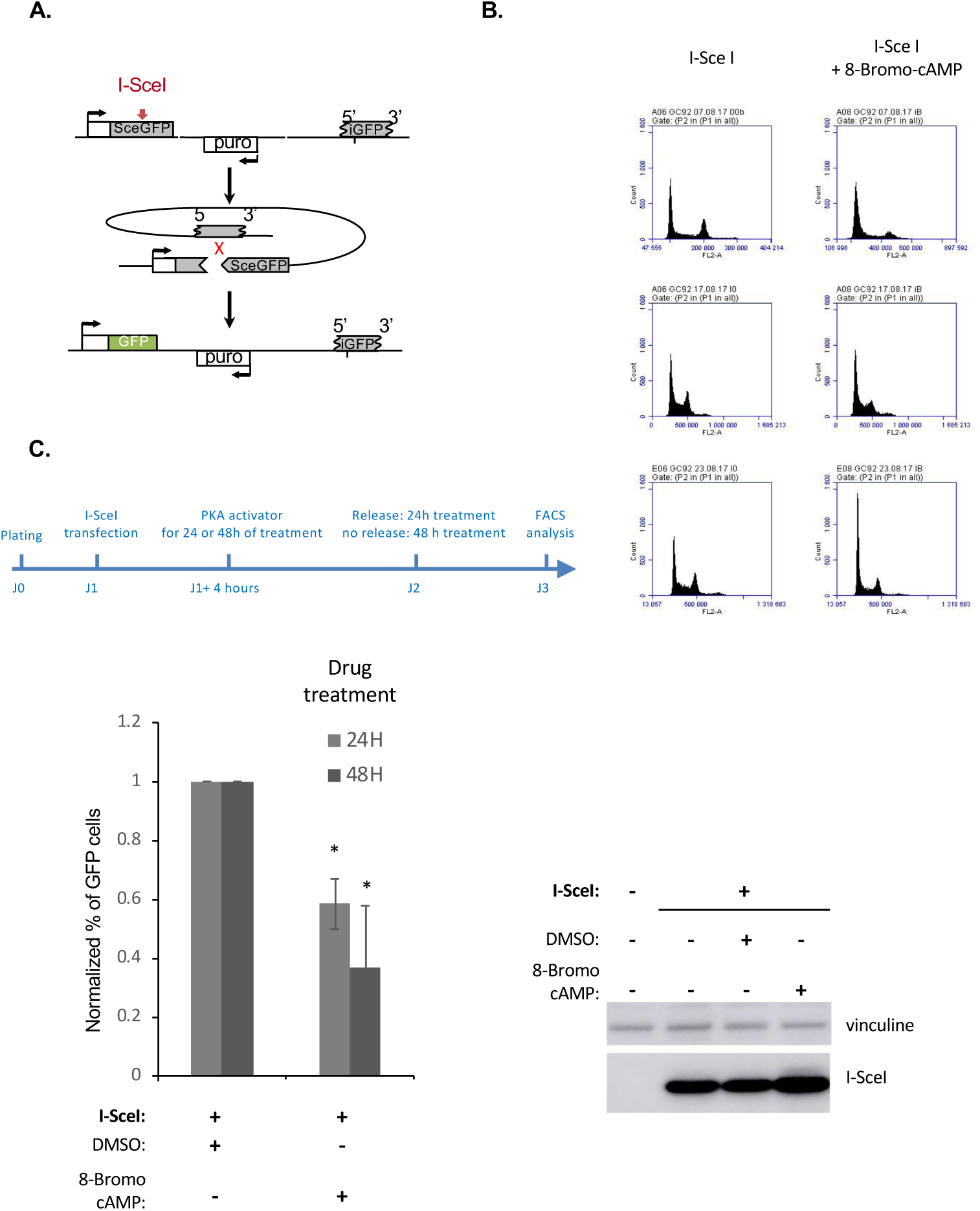
Impact of PKA stimulation on HR. **A.** Strategy and reporter substrate used. Here the DR-GFP substrate is chromosomally integrated into the genome of SV40-transformed human fibroblasts (Dumay et al. 2006). The DR-GFP contains a tandem repeat of two inactive for EGFP encoding genes (upper panel). The 5’ repeat is intereupted by the I-Scei cleavage sequence (18 nt) et the 3’ repaet is trucated both in 5’ and in 3’. None of the mis expressed and the cells are GFP-negative. Expression the I-SceI enzyme generates a cleavage targetted into the 5’ repeat (the SceGFP cassette) ; gene conversion (HR) with the 3’ repeat (iGFP) recreates a functional EGFP leading the fluorescence of cells (Pierce et al. 1999). Recombinant cells can be detected by FACS. **B.** Cell cycle distribution in GC92 cells (human fibroblasts) after I-SceI transfection and PKA activation (8-Bromo-cAmp). The picture shows 3 independent experiments. **C.** Impact of stimulation of PKA by 8-Bromo-cAMP on HR, after I-SceI transfection. Upper panel scheme of the experiment. The values are shown normalized to the control (DMSO) and represent the average ± SEM of at least 3 independent experiments.

Consistently with the data above, stimulation of PKA activity with Bromo-8-cAMP decreased HR efficiency (Figure 6C). The putative small accumulation in the G1 phase might, at least in part, participate to the decrease in HR efficiency, but this seems not really significant.

The engagement of DSB repair toward NHEJ should result in the decrease of competing repair processes such as HR. Collectively, our data are consistent with the stimulation HR after silencing PKA with an siRNA (Adamson et al. 2012) and with the above data on resection and NHEJ.

## Discussion

DSB is a very toxic lesion that can also generates profound genome rearrangements (Bétermier et al. 2014; So et al. 2017). However, DSBs are also employed by cells in physiological mechanisms such as meiosis or the establishment of the immune system. Several repair mechanisms compete for DSB repair. A precise control of DSB repair is essential to select the most appropriate DSB repair mechanism, according to the cell physiology, to maintain genome stability (So et al. 2017; Symington and Gautier 2011). The first step of the choice of the DSB repair pathway is the competition between NHEJ and resection (Rass et al. 2009; Bétermier et al. 2014; So et al. 2017, 2022; Thomas et al. 2023). Resection is necessary to initiate DSB repair by HR. At this essential initial step 53BP1 plays a pivotal role: indeed, it prevents resection, favoring thus NHEJ (So et al. 2017; Symington and Gautier 2011). Here we show that PKA plays a role at this early step of DSB repair. Indeed, we show that PKA physically interacts with 53BP1 facilitating or stabilizing the binding of 53BP1 to damaged DNA. Consistent with these data PKA prevents resection, favoring NHEJ. In a mirror effect, the silencing of PKA, which increases resection as shown here (see Figure 4), has been shown to stimulate HR (Adamson et al. 2012), consistent with the fact that HR is initiated by resection. Coherently, we show here that, reciprocally, stimulating PKA, which stimulates NHEJ (see Figure 5), leads to a decrease in HR (see Figure 6).

Analyzing 53BP1 sequence in phosphosites databases predicted at least 12 potential sites of phosphorylation by PKA in 53BP1. Therefore, whether PKA phosphorylates 53BP1 and identification of the phosphorylated sites of 53BP1 by PKA represents an existing huge challenge for future prospects.

An impact of PKA on NHEJ has been reported, but with contradictory results and conclusions. Noteworthy all these reports show an impact of PKA at late steps of DSB repair, i.e. downstream the choice between NHEJ *versus* resection. A negative impact of cAMP signaling has been described in non-small cell lung cancer (NSLC) cells through the phosphorylation of DNA-PKcs, which plays a central role in NHEJ in mammalian cells, and the recruitment of XRCC4 and ligase IV, i.e. at the last step of NHEJ (Noh and Juhnn 2020). In this report DSB repair was analyzed after IR or on transient episomic plasmids. In both cases this corresponds to hundreds of DSBs; indeed the number of transfected plasmids plasmids can be very high and the numbers of DSB following IR is estimated to 40 DSB/Gy, and the experiments were performed at 2 Gy (Noh and Juhnn 2020). Our reporter substrate was chromosomally integrated, thus with the full chromatin regulation (in contrast with episomic transfected plasmids) and correspond to a unique site. Therefore, it is possible that the impact of PKA on NHEJ might be different on intrachromosomal sequences and/or differs according the number of DNA ends, as reported for MRE11 (Lee and Paull 2004). Contradicting this report, but in agreement with our present data, other reports conclude to a positive impact of PKA on NHEJ. PKA has been shown to favor the entry of DNA-PK into the nucleus in HEK-B2 cells (Huston et al. 2008). PKA activates DNA-PK expression and upregulation of Ligase IV in retinal endothelial cells (Yang et al. 2016; Zhang and Steinle 2013). Notably, the expression of ligase IV is controlled by pCREB, an effector of PKA, in retinal neurocytes (Yang et al. 2016). This is consistent with the activation of CREB phosphorylation after IR, as also shown here. In all these cases PKA favors NHEJ. It has been thus proposed the type of cell tissue might affect the impact of PKA on NHEJ (Noh and Juhnn 2020). Finally, in yeast *Saccharomyces cerevisiae*, the yeast ortholog of PKA, Tpk1, favors NHEJ through the phosphorylation of Nej1, an NHEJ factor (Jessulat et al. 2021). In mammalian cell PKA has been show to phosphorylate Cernunos/XLF, the Nej1 mammalian ortholog that is a co-factor of Ligase IV, thus acting at the last step of NHEJ. Here we show that PKA physically interacts with 53BP1, preventing resection, i.e. acting at the first step of DSB repair, orienting the choice of the repair mechanisms. Collectively, these data underline the high complexity of the regulation of DSB repair by PKA that can act both at early and late steps of NHEJ. The roles of PKA on NHEJ are summarized in Figure 7.

**Figure 7:**
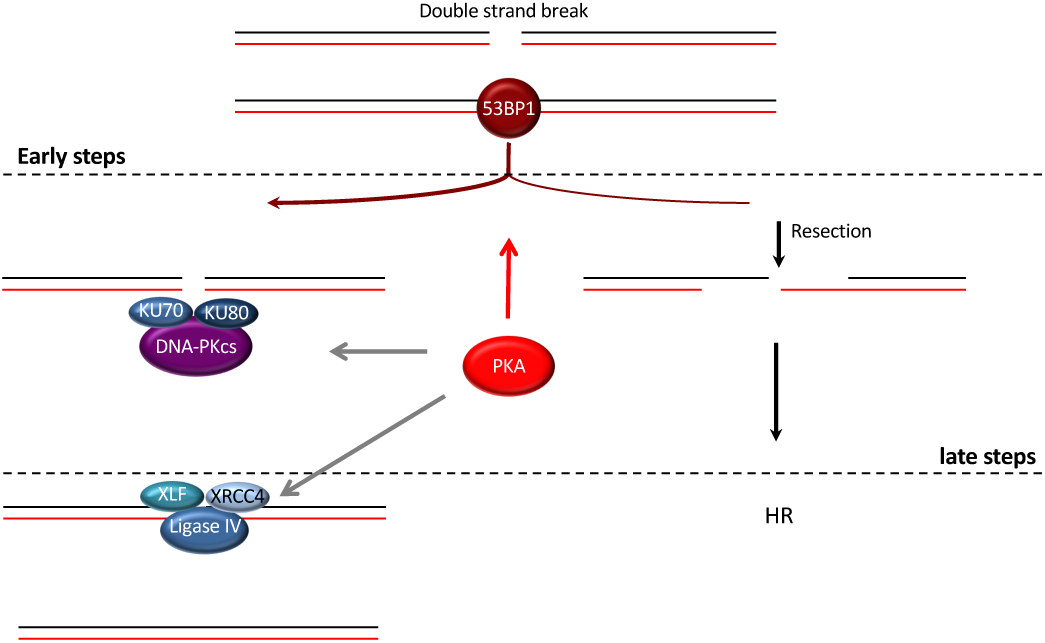
The different role of PKA on NHEJ. At the earliest step of DSB repair, PKA favors and/or stabilizes the loading of 53BP1 on the DSBs, impairing resection and fostering NHEJ. Resection repression leads to the decreased HR efficiency. However PKA can also act at later steps of NHEJ, through the nuclear translocation of DNA-PK and the expression of NHEJ factors including the ligase IV and its cofactors (XRCC4 and XLF/Cernunos).

We show that stimulating PKA activity decreases HR. This is coherent with the fact that this stimulates in parallel NHEJ. Therefore, stimulation of PKA activity, by modifying the balance NHEJ *versus* HR in favor of NHEJ, should lead to a decrease in HR.

These data potentially open new avenues for future therapeutic strategies. Indeed, since NHEJ plays an important role in resistance to ionizing radiation, decreasing NHEJ efficiency, though PKA inhibition, might increase the efficiency of radiation therapy. Moreover, HR deficient cells are sensitive to PARP inhibitors trough a synthetic lethality process (Farmer et al. 2005; Bryant et al. 2005). Unfortunately, most tumors are proficient for HR. Therefore, inhibition of HR might make potentially any tumor eligible to PARP inhibitors therapy. Our data reveal that stimulation of PKA activity can alter HR. However, increasing NHEJ by PKA stimulation might counteract the sensibility to PARP inhibitors resulting from HR inhibition. Therefore, these characteristics of PKA on DSB repair might allow a combination of different alternative strategies that deserve to be address in future prospects, giving the opportunity to adapt potential treatments.

PKA should be considered as an actual actor of DDR. Indeed, our data reveal PKA an important novel actor at initial step of DSB repair, controlling the choice of the DSB repair process, which might have impact on genome stability maintenance and open potential enticing novel therapeutic strategies. This might confer a role to PKA in very precise regulation(s), allowing selecting the most appropriate DNA repair system according to different cellular and tissues parameters, in order to efficiently maintain genome stability.

## Acknolewledgements

This work was supported by grants from the Institut National du Cancer (PLBIO21-072), La Ligue Contre Le Cancer (ARN therapeutiques), and ITMO Cancer (PCSI 2022), Fondation ARC (ARCPJA2022060005157).

